# Copper Sulfide Nanoparticle–Mediated Photothermolysis Induces Immunogenic Cell Death in Ovarian Cancer Cells

**DOI:** 10.1101/2025.07.08.662637

**Authors:** Dan Li, Sixiang Shi, Qizhen Cao, Anil K. Sood, Jeffrey J. Molldrem, Qing Ma, Chun Li

**Affiliations:** Departments of Hematopoietic Biology and Malignancy, The University of Texas MD Anderson Cancer Center, Houston, Texas 77054, United States; Departments of Cancer Systems Imaging, The University of Texas MD Anderson Cancer Center, Houston, Texas 77054, United States; Departments of Gynecologic Oncology and Reproductive Medicine, The University of Texas MD Anderson Cancer Center, Houston, Texas 77054, United States

**Keywords:** Photothermolysis, CuS nanoparticles, Dendritic cells, T cells

## Abstract

Immunotherapy has only limited efficacy against ovarian cancer because of dysfunctional T cells in an immunosuppressive tumor microenvironment. Selective photothermolysis is a technique in which short pulsed laser is used to ablate tissue in targeted regions in a precise spatiotemporally controlled manner. This technique allows targeted tissue ablation without damage to surrounding tissue. In the current study, we demonstrated that ovarian cancer cells treated with photothermolysis mediated by near-infrared-absorbing CuS nanoparticles and 15-ns laser pulses were ingested by human monocyte-derived dendritic cells (MoDCs) more efficiently than were ovarian cancer cells treated with ionizing radiation. Moreover, ingestion of ovarian cancer cells treated with CuS NP-mediated photothermolysis promoted MoDC activation. Immature MoDCs that had been exposed to photothermolysis-treated cancer cells caused greater clonal expansion of co-cultured human CD4^+^ and CD8^+^ T cells than did immature MoDCs that had been exposed to ionizing radiation–treated tumor cells. These data indicate that photothermolysis may be used to selectively target and destroy tumor cells and cause them to release antigens and danger signals, which can be recognized by DCs to activate T cells.

## INTRODUCTION

Local tumor ablation techniques including laser-mediated ablation have been tested to expose tumor-associated antigens and enhance antigen presentation to T cells **[1-3]**. Among these techniques, laser-induced tumor ablation is unique in that it offers high selectivity, which can be increased by introducing near-infrared (NIR) light–absorbing nanoparticles (NPs) that are selectively delivered to tumor cells **[4-6]**. Conventional laser-induced tumor ablation uses continuous-wave laser to induce necrotic cell death at temperature greater than 50°C **[2, 3]**. Alternatively, short nanosecond pulsed-wave (PW) laser produces a photomechanical force to kill cells. This phenomenon is termed photothermolysis **[7, 8]**. In photothermolysis, the photoenergy is converted to thermomechanical energy when laser beam is absorbed **[9, 10]**. When treatment conditions (i.e. power density and duration of treatment) are equal, PW laser has a much greater tissue penetration than continuous-wave laser **[11]**. Our computer simulation showed that the thermal energy generated from a point source by a single 15-ns laser pulse is confined within a time frame of 10 ns and a spatial frame of less than 0.1 μm **[12]**, suggesting that heat does not dissipate beyond a few cells when treatment duration is limited to seconds. Therefore, PW laser– mediated cell death can be controlled in a highly spatiotemporally confined fashion. This is relevant for a disease like ovarian cancer where damage to surrounding normal tissues during surgery, such as the bowel, can cause potential risks and complications. It is important that innovative treatments are developed to minimize such risks while effectively eliminating residual tumor cells.

Before photothermolysis can be translated to the clinic as a new cancer treatment modality, two fundamental issues must be addressed. First, it must be demonstrated that photothermolysis can induce a systemic response, i.e., that cell death induced by photothermolysis is capable of activating dendritic cells (DCs) and T cells. Second, suitable NIR light–absorbing materials must be identified and developed to mediate efficient photothermolysis. NIR light–absorbing materials should be biodegradable and biocompatible to minimize potential toxic side effects. It is also desirable to have an absorption spectrum that peaks at 1064 nm to match laser beam generated by most clinically used PW laser systems. NIR light–absorbing Au-based NPs (nanorods, nanoshells) can remain in the body for months, which raises safety concerns **[13]**. Moreover, tunable Ti-Sapphire laser, which delivers pulses in the range of 700-960 nm, is only 20% as powerful as 1064-nm Q-switched Nd:YAG laser.

In the study reported here, we demonstrated that tumor cells treated with biodegradable polyethylene glycol (PEG)-coated copper sulfide NPs (PEG-CuS NPs) and 15-ns pulsed laser at 1064 nm can activate DCs and T cells *in vitro*, supporting a treatment strategy in which DCs are exposed to photothermolysis-treated tumor cells to induce potent antigen-specific immune responses.

## RESULTS

### Synthesis and characterizations of PEG-CuS NPs

PEG-CuS NPs were generated according to a previously reported method **[14, 15]**. The resulting NPs had a number average diameter of 11 nm and a volume average diameter of 26 nm (**Fig. 1A**). The size of the NPs was also confirmed by transmission electron microscopy (**Fig. 1B**). PEG-CuS NPs were irregular in shape. PEG-CuS NPs displayed peak absorption at 1058 nm (**Fig. 1C**), which matches the wavelength of the Nd:YAG laser at 1064 nm.

**Fig. 1.**
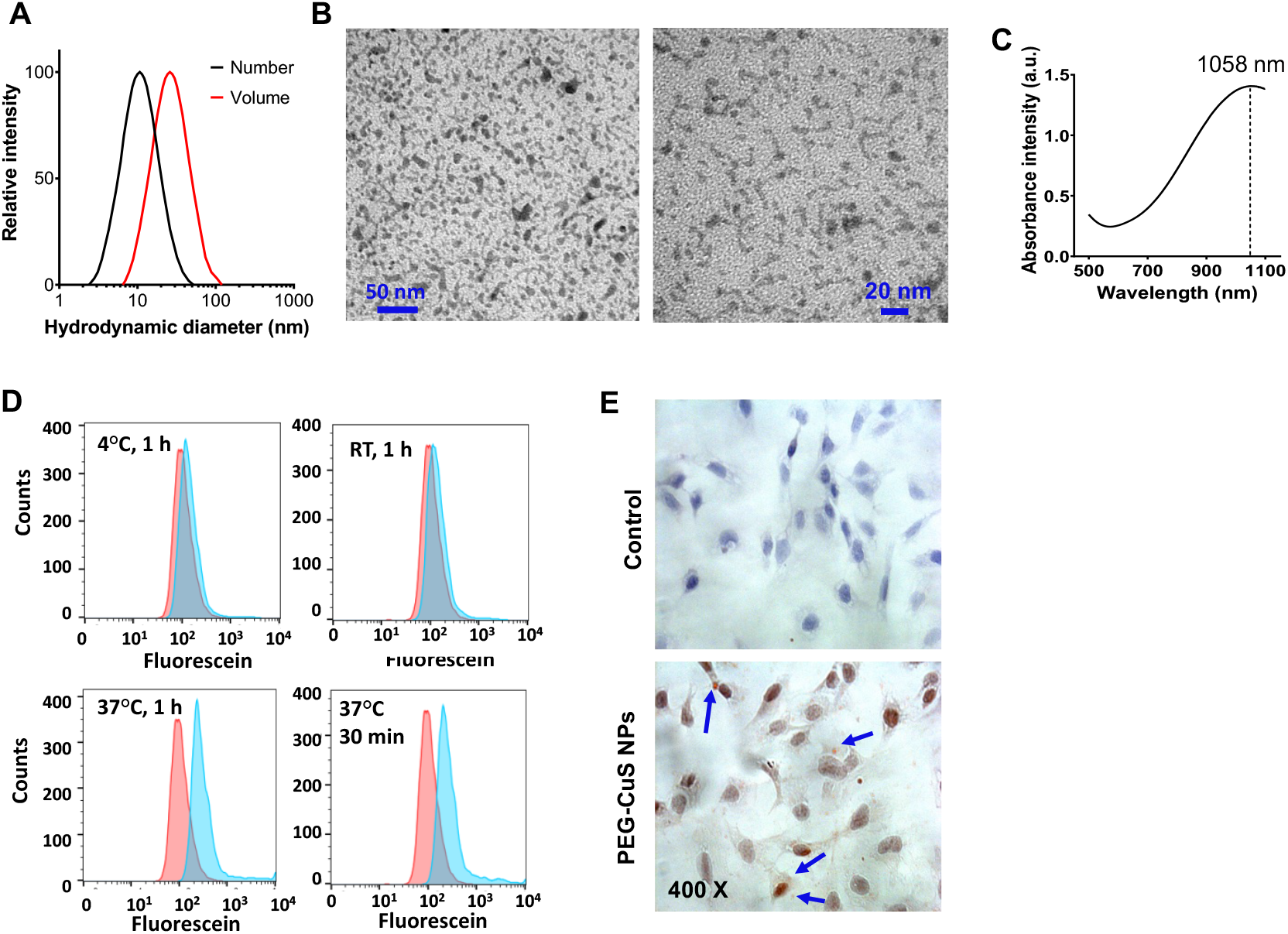
Synthesis and characterization of PEG-CuS NPs. (**A**) Hydrodynamic sizes of PEG-CuS NPs measured by dynamic light scattering. (**B**) Transmission electron microscopy microphotograph of PEG-CuS NPs (original magnification ×300,000). (**C**) Absorbance profile of PEG-CuS NPs measured by ultraviolet-visible spectrometry. (**D**) Flow cytometry histograms of SKOV3 cells treated with carboxyfluorescein succinimidyl ester–labeled PEG-CuS NPs at different temperatures for 30 min and 1 h. Red: untreated SKOV3 cells; blue: SKOV3 cells treated with fluorescence-labeled PEG-CuS NPs. RT, room temperature. (**E**) Representative microphotographs of PBS-treated SKOV3 cells (top; control) and SKOV3 cells incubated with PEG-CuS NPs (bottom) stained with rhodanine and hematoxylin. Dark red color (arrows) indicated the presence of rhodanine-stained Cu element and confirmed NP internalization. Cells were incubated at 37°C for 1 h and visualized under a microscope at the original magnification of ×400.

PEG-CuS NPs were taken up by SKOV3 human ovarian cancer cells when incubated at 37°C. Flow cytometry study of SKOV3 cells treated with NHS-fluorescein (carboxyfluorescein succinimidyl ester)–labeled CuS NPs showed uptake of the PEG-CuS NPs in the tumor cells when they were incubated at 37°C for 30 min or 1 h but not when they were incubated at 4°C or room temperature (**Fig. 1D**). These data suggest that cellular uptake of PEG-CuS NPs was mediated by active transport processes such as nonspecific endocytosis of NPs. Intracellular deposition of PEG-CuS NPs in SKOV3 cells was confirmed by the presence of rhodanine-stained Cu element revealed under a photomicroscope (**Fig. 1E**).

### Photothermolysis mediated by PEG-CuS NPs can increase ingestion of tumor cells by MoDCs and activation of MoDCs

Immunogenic cell death is accompanied by rapid release of tumor-associated antigens and damage-associated molecular patterns, which can induce antitumor immunity if the tumor-associated antigens are efficiently taken up and processed by DCs and presented to naïve T cells **[16]**. We compared DC phagocytosis of photothermolysis-treated tumor cells and tumor cells treated with ionizing radiation (IR) delivered at 3000 cGy. The IR dose of 3000 cGy was chosen based on a report by Huang et al. [**17**] that minimal DC cell death was observed following *in vitro* treatment with 3000 cGy. On the other hand, irradiation of SKOV3 cells at 600 cGy caused 50% apoptotic cell death [**18**], and at a dose of 800 cGy there was less than 10 percent survival cells in an in *vitro* colony formation assay [**19**]. While only 2% to 3% of IR-treated SKOV3 tumor cells were ingested by human monocyte-derived DCs (MoDCs) **[20]**, 12% to 23% of SKOV3 cells treated with PEG-CuS NPs and 15-ns pulsed laser were taken up by MoDCs (**Fig. 2**). Thus, photothermolysis-treated tumor cells were efficiently ingested by MoDCs.

**Fig. 2.**
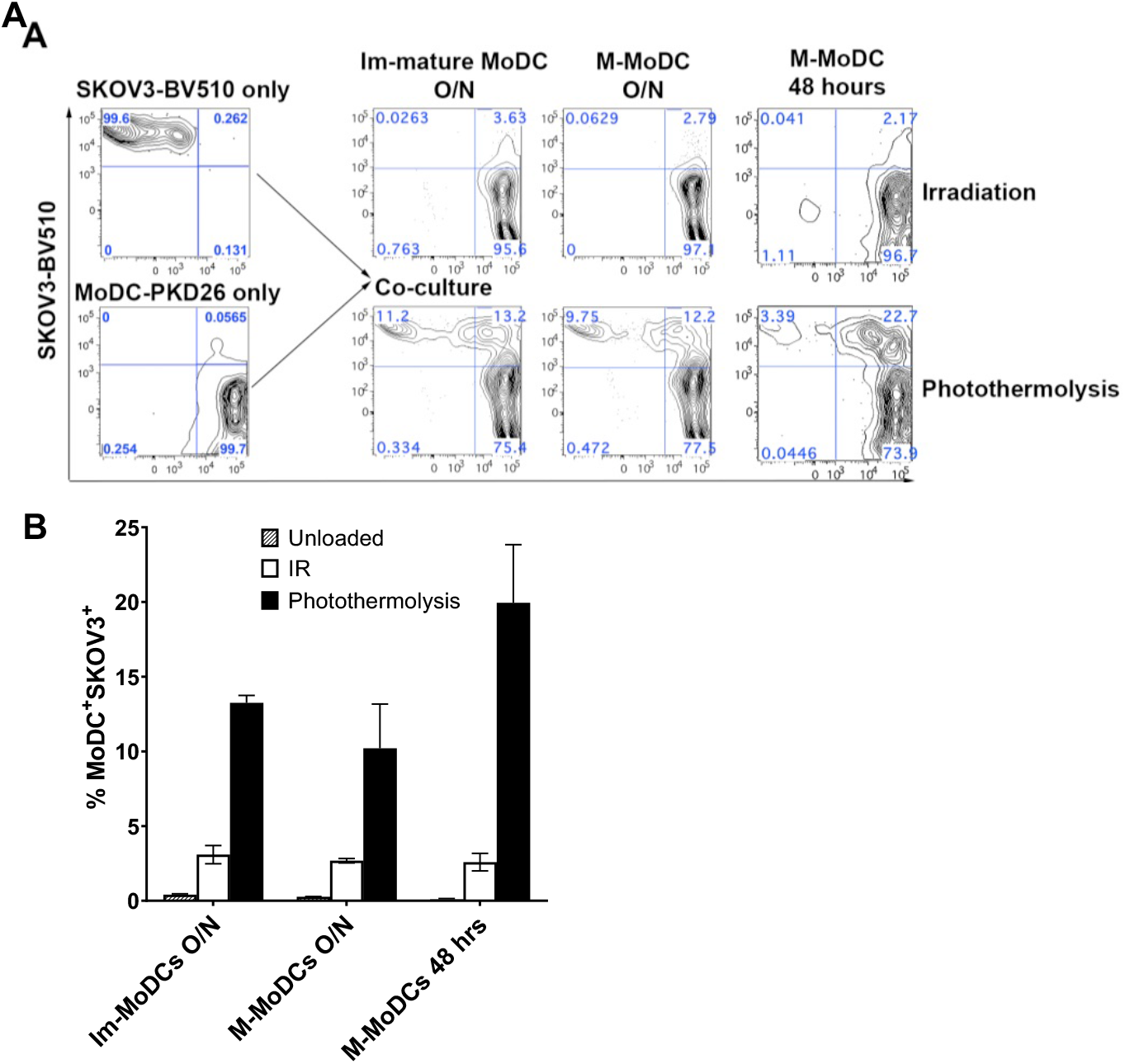
Tumor cells treated by photothermolysis were efficiently ingested by MoDCs. (**A**) Representative flow cytometry dot plots showing the percentage of treated SKOV3 tumor cells ingested by MoDCs. MoDCs were labeled with PKH26 dye; tumor cells were labeled with BV510 dye. Cells were exposed to PEG-CuS NPs (100 μg/mL) at 37°C for 20 min and washed extensively with PBS before treatments. IR was delivered at 3000 cGy. Photothermolysis was instituted by irradiation with 15-ns laser pulses at 1064 nm (0.25 mJ/cm^2^, 60 sec). (**B**) Quantification of ingested SKOV3 cells treated by either IR or photothermolysis. Immature MoDCs without addition of tumor cells were used as a control. Data are expressed as mean ± SD (n = 2). MoDCs were incubated with treated tumor cells overnight (O/N) or for 48 h. Im-MoDCs, immature MoDCs; M-MoDCs, mature MoDCs.

Ingestion of SKOV3 cells dying as a result of photothermolysis altered the phenotypes of MoDCs, as indicated by significantly greater expression levels of CCR7 (homing marker) and CD14 than were observed in MoDCs incubated with IR-treated SKOV3 cells (p<0.05) (**Fig. 3A, 3B**). CD14 binds to damage associated molecular patterns (DAMPs) and toll-like receptors to engage in activation of innate immune system [**21**]. At 48 h, the majority of the MoDCs expressed CD83 (maturation marker) (>83%). At such as high baseline level, it was difficult to differentiate expression levels among different treatment groups (**Fig. 3C**). Both IR-treated and photothermolysis-treated SKOV3 cells resulted in increased CD83 expression on MoDCs after 48 h of incubation.

**Fig. 3.**
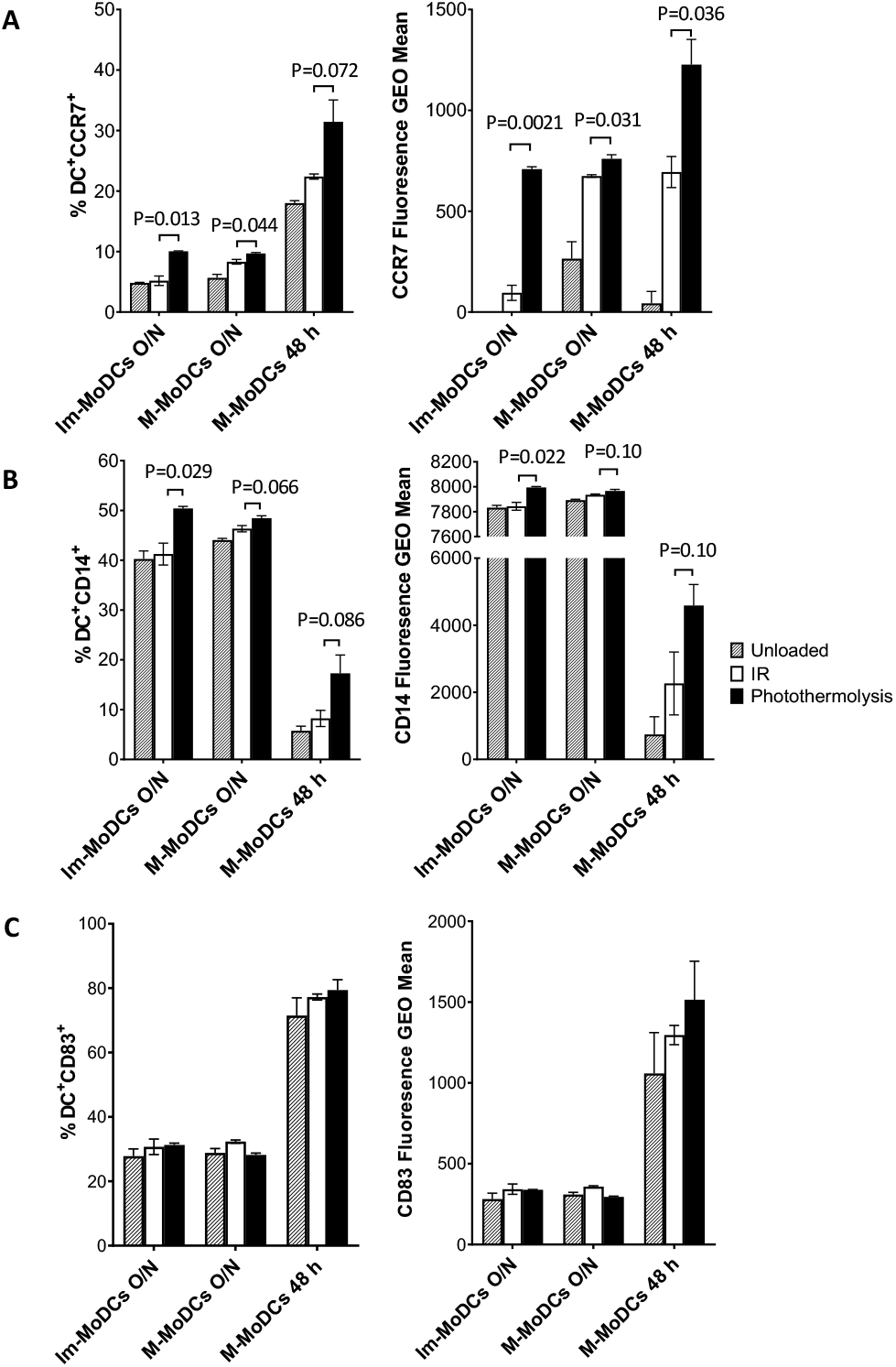
Ingestion of ovarian tumor cells treated with CuS NP-mediated photothermolysis promoted DC activation. Expression of (**A**) CCR7, (**B**) CD14, and (**C**) CD83 on MoDCs exposed to SKOV3 tumor cells. Tumor cells were incubated with PEG-CuS NPs (100 μg/mL) for 20 min. After extensive washing, cells were treated with either IR (3000 cGy) or photothermolysis (15-ns pulses, 250 mJ/cm^2^, 600 pulses) and then co-cultured with MoDCs overnight (O/N) or for 48 h before flow cytometry analysis. Data are expressed as mean ± SD (n = 2). Im-MoDCs, immature MoDCs; M-MoDCs, mature MoDCs; Fluorescence GEO Mean, geometric mean fluorescence intensity.

### Activated MoDCs can promote T-cell proliferation

To further determine the activity of MoDCs, carboxyfluoroscein succinimidyl ester (CFSE)-labeled peripheral blood mononuclear cells (PBMCs) from healthy donors were co-cultured with MoDCs activated by ingestion of SKOV3 cells dying as a result of photothermolysis. T-cell proliferation and activation were measured on day 5 using FACS. As shown in **Figure 4**, loading of immature MoDCs with photothermolysis-treated SKOV3 cells led to a significant increase of CD4^+^ T-cell proliferation compared to the level in MoDCs loaded with IR-treated SKOV3 cells (p<0.05). Although the p value for the percentage of proliferating CD8+ T cell as not statistically significant between the IR and the photothermolysis treatment group (p=0.78), the total number of live CD8+ T cells showed significantly difference between IR treatment and photothermolysis (p=0.0077) (**Fig. 4**). Thus, ingestion of photothermolysis-treated tumor cells can activate MoDCs and promote T-cell proliferation, generating a potent immune response.

**Fig. 4.**
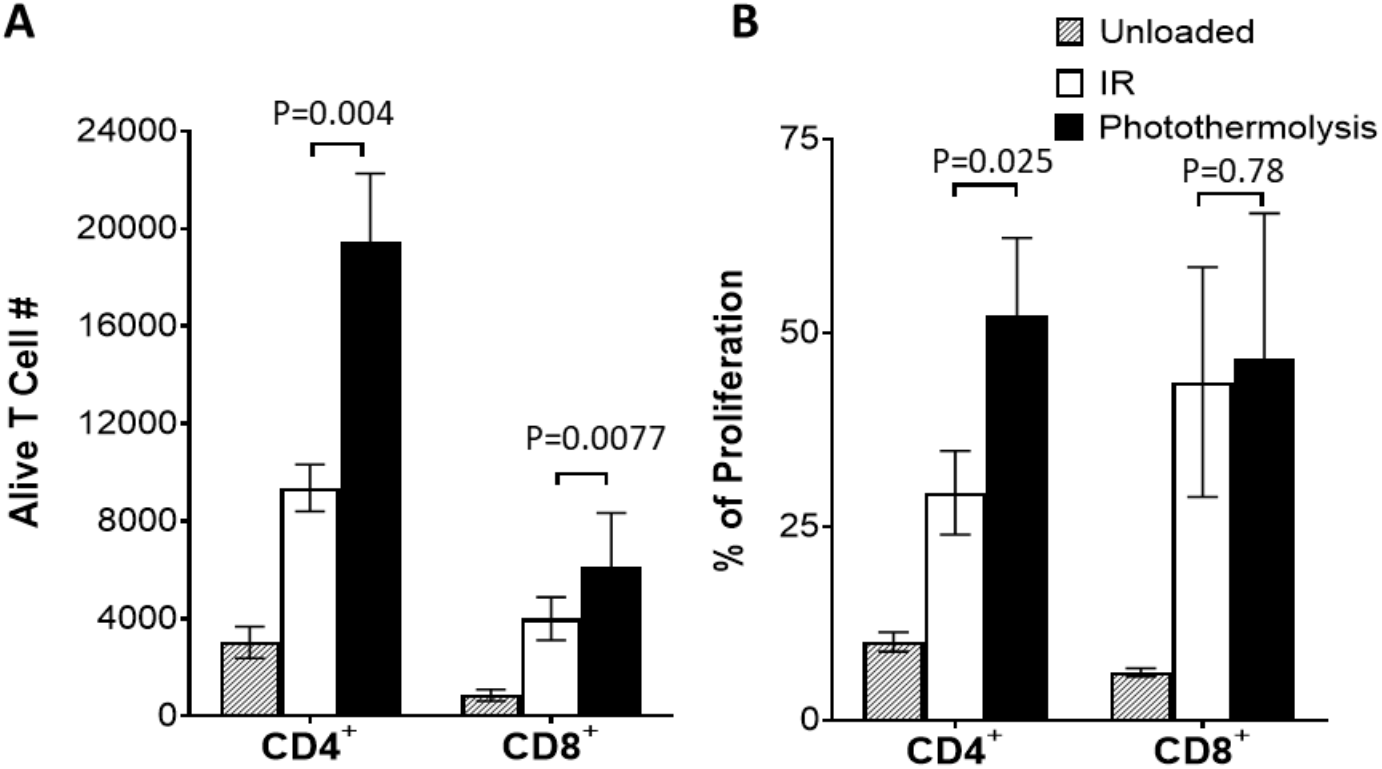
Clonal expansion of CD4^+^ and CD8^+^ T cells after co-culture with Im-MoDCs that had been exposed to IR-treated or photothermolysis-treated SKOV3 tumor cells. (**A**) Numbers of CD4^+^ and CD8^+^ T cells alive and (**B**) percentages of CD4^+^ and CD8^+^ T cells proliferating after 5-day incubation with Im-MoDCs. Bar graphs represent the mean ± SD; n = 3. CFSE-labeled PBMCs were co-cultured for 5 days with immature MoDCs loaded with SKOV3 cells treated with either IR or photothermolysis. CFSE-labeled PBMCs co-cultured with unloaded MoDCs were used as a negative control. Flow cytometry was used to detect CFSE dilution as a measure of T-cell proliferation. CD4^+^ T cells were identified with PerCP-5.5-conjugated anti-CD4, and CD8^+^ T cells were identified with PE-Cy7-conjugated anti-CD8.

## DISCUSSION

In this study, we demonstrated that human DCs loaded with tumor cells dying as a result of photothermolysis facilitated *ex vivo* uptake of tumor-associated antigens and effective processing and stimulation of T cells, which may lead to enhanced immunogenicity and effective DC therapy.

One of the primary therapeutic challenges in ovarian cancer is widespread metastases at the time of diagnosis, which can be difficult to surgically resect **[22]**. For this reason, new treatments are urgently needed. Previously, we showed that PEG-CuS NP–mediated photothermolysis in combination with anti-PD-1 checkpoint inhibitor and the toll-like receptor 9 agonist PG-CpG could induce systemic antitumor immunity in a poorly immunogenic murine ovarian cancer model **[15]**. However, clinical translation of this approach is limited by limited efficiency of NP delivery to tumors after systemic administration and the need for regulatory approval for systemic administration of PEG-CuS NPs, which is a costly and lengthy process.

DCs are professional antigen-presenting cells. DC-based immunotherapy is a particularly attractive option because DCs can be exposed to dying tumor cells treated by photothermolysis *ex vivo*. DC-based vaccines have been investigated in numerous clinical trials and proven to have low toxicity and the potential to induce long-term immunological memory **[23]**. One example demonstrating feasibility and clinical translatability of DC-based therapy is the approval of Sipuleucel-T for the treatment of castrate-resistant metastatic prostate cancer. Sipuleucel-T is a product made by *ex vivo* exposure of a patient’s own immature antigen-presenting cells collected by leukapheresis to a fusion protein composed of human granulocyte-macrophage colony-stimulating factor (GM-CSF) and prostatic acid phosphatase, a prostate cancer–specific antigen **[24]**. Sipuleucel-T has demonstrated a survival benefit and a high benefit-risk ratio and represents a first-in-class autologous DC therapy.

To date, DC-based vaccines have not fulfilled their promise for the treatment of ovarian cancer. To improve the efficacy of DC-based immunotherapy, it is imperative that DCs be optimized against individual tumors by enhancing their maturation status and migratory potential. In the current study, we demonstrated for the first time that PEG-CuS NP–induced photothermolysis of human SKOV3 ovarian tumor cells could enhance ingestion of tumor cells by human MoDCs. Importantly, photothermolysis-treated tumor cells induced more potent activation of MoDCs and human T-cell proliferation than did IR-treated tumor cells. We used nontargeted PEG-CuS NPs because the NPs could be taken up by SKOV3 ovarian cancer cells. It is expected that CuS NPs targeted to receptors expressed on the surface of ovarian cancer cells could further increase cellular uptake of CuS NPs via receptor-mediated uptake.

Because of the heterogeneity of ovarian cancer and other solid tumors, targeting multiple unique antigens restricted to individual tumors is the most important parameter in advancing DC-based immunotherapy **[25]**. Whole tumor cell–based cancer vaccines may present not only antigens commonly associated with the tumor type but also patient-specific neoantigens and may therefore reduce the potential for immunological tolerance and be robustly immunogenic, because such approaches have the potential to simultaneously present both shared tumor-associated antigens and private mutated neoantigens. A recent pilot clinical study showed that an oxidized whole-tumor lysate DC vaccine was safe and elicited broad antitumor immunity **[26]**.

At present, the mechanisms of enhanced phagocytosis of short-laser pulse-treated SKOV3 cells by MoDC are not clear. Previously, we have observed that cell death accompanied by blebbing of cell membrane, rounded morphology, and detachment from microplate surfaces within a minute after one shot of short pulsed laser [**12**]. This process was extremely rapid, suggesting that cell death is mostly likely caused by cell membrane damage rather than a form of programmed cell death (i.e., apoptosis or ferroptosis). Ogawa et al. [**2**7] reported NIR photoimmunotherapy induced by IR700 dye-anti-HER2 antibody conjugate using continuous wave diode laser at 670–690 nm. Laser treatment induced immunogenic cell death and DC maturation. Alzeibak et al. [**28**] reviewed recent progress on NIR light induced immunogenic cell death, in particular in the context of photodynamic therapy. We emphasize the fact that short-pulsed (15-ns) laser differs from continuous-wave laser in that the thermo-mechanical energy is confined in the former case while thermo-energy is dissipated to the environment in the latter case. Clearly, more studies are needed to establish the specificity and usefulness of photothermolysis in inducing immunogenic cell death compared to photodynamic therapy and photothermotherapy that use continuous-wave laser beam.

Several limitations of this study are noted. First, *in vivo* studies are needed to show that MoDCs incubated *ex vivo* with photothermolysis-treated tumor cells migrate efficiently to the lymph nodes and are effective in preventing ovarian cancer after tumor cell challenge in the vaccine setting or eliminating ovarian cancer cells in the therapy setting. Second, it is likely that DC activity may be further enhanced by providing cytokine cocktails and costimulatory factors (e.g., TLR ligand, GM-CSF, IL-2), and further studies are needed to optimize the treatment protocol to further boost activation and mobility of DCs. Lastly, clinical translation of such a method is needed. The *ex vivo* DC activation method described in this study will expedite future clinical translation. Third, the data on temperature change at the cellular level was not measured at the due to limitation of available experimental tool. However, previous computer simulation with one pulse of 15-ns laser showed that temperature elevation is confined to a radius of 0.1 μm and a time scale of 100 ns [**12**], which suggest that temperature increase would be confined to individual cells that have taken up NIR light absorbing nanoparticles and would not spread to surrounding normal tissue. In fact, the Nd:YAG pulsed 15-ns laser used in our experiments is widely employed in aesthetic and medical applications.

## MATERIALS AND METHODS

### Materials

Copper (II) chloride (CuCl_2_), sodium sulfide (Na_2_S·9H_2_O), poly(ethylene glycol) methyl ether thiol (mPEG-SH; molecular weight: 5000), folic acid, and 1-ethyl-3-(3-dimethylaminopropyl)carbodiimide were purchased from Sigma-Aldrich (St. Louis, MO). Heterobifunctional PEGylation reagent thiol PEG amine (HS-PEG-NH_2_; molecular weight: 5000) was purchased from Creative PEGworks (Chapel Hill, NC). All other chemicals were purchased from ThermoFisher Scientific (Waltham, MA). PD-10 desalting columns were purchased from GE Healthcare Life Sciences (Pittsburgh, PA). Amicon ultra-15 centrifugal filters were purchased from EMD Millipore (Burlington, MA). All other antibodies were purchased from either BD Biosciences (San Jose, CA) or BioLegend (San Diego, CA).

### Synthesis and characterizations of PEG-CuS NPs and fluorescein-PEG-CuS NPs

The general procedure for synthesis of PEG-CuS NPs was modified from our previous report **[14]**. In brief, 6 mg of mPEG-SH and 4 mg of HS-PEG-NH_2_ were added into 10 mL of deionized water (18 MΩ), and the PEG solution was vigorously stirred at room temperature for 5 min. CuCl_2_ (5 μmol) and Na_2_S (10 μmol) were sequentially added into the PEG solution under stirring. Subsequently, the reaction mixture was moved to a 95ºC oil bath and heated for 15 min under stirring until a dark green solution was obtained. The resulting product, NH_2_-PEG-CuS, was moved onto an ice bath until it cooled to room temperature. Ultrafiltration was performed using Amicon ultra-15 centrifugal filters (molecular weight: 30,000) at 5000 × g for 12 min to remove unreacted CuCl_2_, Na_2_S, and HS-PEG-NH_2_. The purification process was performed a total of three times, and 1 mg of mPEG-SH was added into the sample for each purification to prevent aggregation. The final product was stored at 4ºC until use.

NHS-fluorescein (5/6-carboxyfluorescein succinimidyl ester) was conjugated onto NH_2_-PEG-CuS NPs for *in vitro* flow cytometry analysis. The conjugation was conducted with an amine-to-NHS-fluorescein molar ratio of 1:25 at pH 8 at room temperature for 2 h. The resulting product, fluorescein-PEG-CuS NPs, was purified with a PD-10 desalting column to remove free dye. Successful conjugation of fluorescein to the amine-terminus of PEG in PEG-CuS was confirmed by fluorescence spectrophotometry.

### Measurement of NPs

NP size was characterized by dynamic light scattering (Brookhaven Instruments Corporation, Holtsville, NY) and transmission electron microscopy (JOEL JEM-1010 microscope; JOEL USA, Inc., Peabody, MA). The absorbance spectrum was acquired with an ultraviolet-visible-NIR spectrophotometer (model DU 800; Beckman Coulter, Brea, CA).

### Cancer cell culture and NP uptake

SKOV3 human ovarian cancer cells were obtained from American Type Culture Collection (Manassas, VA) and cultured in RPMI 1640 cell culture medium. Cells were digested with 0.25% (w/v) trypsin for *in vitro* experiments when they reached ∼80% confluence.

Uptake of fluorescent labeled PEG-CuS NPs in SKOV3 cells was analyzed by flow cytometry (FACSCalibur; Becton-Dickinson, San Jose, CA). In brief, 0.5 × 10^6^ SKOV3 cells were harvested and incubated with fluorescein-PEG-CuS NPs (5 μg/mL) at 4ºC, 25ºC, and 37ºC for 1 h. The cells were resuspended with cold PBS containing 1% bovine serum albumin after washing steps, and then subjected to flow cytometry analysis. Untreated cells were used as a control.

For photomicrographic study, SKOV3 cells were incubated with PEG-CuS NPs (50 μg/mL) for 2 h at room temperature. Subsequently, the cells were stained with p-dimethylaminobenzylinene-rhodanine (rhodanine) (Newcomer Supply, Middleton, WI) to reveal CuS, which exhibited a dark red color. The cells were counterstained with hematoxylin. Cells were observed using a Zeiss Axio Observer.Z1 fluorescence microscope.

### Photothermolysis and IR

SKOV3 ovarian cancer cells were seeded onto a 96-well plate (10,000 cells per well) 1 day before the experiment. Cells were washed three times with Hank’s balanced salt solution (Sigma-Aldrich), incubated with PEG-CuS NPs at a concentration of 100 μg/mL in McCoy’s 5a Medium at 37°C for 20 min, and then extensively washed. For photothermolysis, a Nd:YAG laser (1064 nm) was used. The cells were exposed to a 15-ns laser beam for 60 s (0.25 mJ/cm^2^, 600 pulses). For treatment with IR, the cells were irradiated at a dose of 3000 cGy with a Mark I-30 ^137^Cs irradiator (JL Shepherd and Associates, San Fernando, CA). After treatment with photothermolysis or IR, the cells were resuspended in cell culture medium (RPMI 1640) for further experiments.

### Generation of MoDCs

MoDCs were generated according to our published protocol **[20]** from monocytes isolated from human PBMCs. Briefly, buffy coats from healthy human donors were obtained from the blood bank of MD Anderson Cancer Center. PBMCs were isolated from buffy coats by separation with Histopaque-1077 (Sigma-Aldrich). Then 1 × 10^7^ PBMCs were loaded in each well of a six-well plate suspended in 3 mL of macrophage-serum free media (M-SFM) (ThermoFisher) and incubated to adhere for 1.5 h. Nonadherent cells were then removed, and the remaining adherent monocytes were cultured in M-SFM supplemented with 100 ng/mL GM-CSF (Sanofi, Bridgewater, NJ) and 100 ng/mL IL-4 (BD Biosciences). Immature MoDCs were then collected from plates after 5 days of incubation at 37°C. Further MoDC maturation was induced by co-culturing immature MoDCs with SKOV3 cells in M-SFM for 1 h and then in M-SFM supplemented with 10 ng/mL IL-1β, 10 ng/mL IL-6, 10 ng/mL TNFα (BD Biosciences), and 10 ng/mL PGE2 (Sigma-Aldrich) for either 12 h or 48 h at 37°C.

### Ingestion of SKOV3 cells by MoDCs

MoDCs were labeled with PKH26 fluorescence dye using the PKH26 Red Fluorescence Cell Linker Mini Kit (Sigma-Aldrich), and SKOV3 cells were labeled with BV510 fluorescence dye using the CytoPainter Live Cell Labelling kit (Abcam, Waltham, MA). SKOV3 cells were exposed to PEG-CuS (100 μg/mL) for 20 min and extensively washed. PKH26-labeled immature or mature MoDCs were co-cultured with BV510-labeled SKOV3 cells treated with either 15-ns laser pulses (1.4 W/cm^2^, 60 sec; 0.25 mJ/cm^2^ for 600 pulses) or 3000 cGy IR either overnight (12 h) or for 48 h at 37°C. Unloaded MoDCs were used as a negative control. Ingestion of SKOV3 cells by MoDCs was analyzed by flow cytometry using a FACSCanto II flow cytometer (BD Biosciences), and the resulting data were analyzed using FlowJo software.

### Immune-phenotyping of MoDCs

MoDC phenotype was determined by flow cytometry analysis 48 h after co-culture with photothermolysis- or IR-treated SKOV3 cells. Cells were labeled with fluorescence-conjugated antibodies, including Alexa 488–conjugated anti-CCR7 (clone G043H7), Pacific blue– conjugated anti-CD14 (clone HCD14), and PerCP5.5–conjugated anti-CD83 (clone HB15e) (BioLegend, San Diego, CA). Unloaded MoDCs were used as a negative control. The expression level of each surface marker on MoDC was detected by flow cytometry using a FACSCanto II flow cytometer, and the resulting data were analyzed using FlowJo software.

### T-cell proliferation assay

Human PBMCs from healthy donors were labeled with 1 μM CFSE (ThermoFisher) and then co-cultured with allogeneic immature MoDCs loaded with either photothermolysis-treated or IR-treated SKOV3 cells for 5 days. CFSE-labeled PBMCs co-cultured with unloaded MoDCs were used as a negative control. The cells were harvested and stained with surface markers, and flow cytometry was used to detect CFSE dilution as a measure of T-cell proliferation. CD4^+^ T cells were identified with APC-conjugated anti-CD3 (clone HIT3α) and PerCP-5.5-conjugated anti-CD4 antibodies (clone SK3), while CD8^+^ T cells were identified with APC-conjugated anti-CD3 and PE-Cy7-conjugated anti-CD8 antibodies (clone HIT8α) (BD Biosciences) using a FACSCanto II flow cytometer, and the resulting data were analyzed using FlowJo software.

### Statistics

Statistical analysis was performed using multiple unpaired t tests with a GraphPad Prism software version 9.0.0 (SPSS Standard version 13.0, SPSS Inc.). Data were expressed as mean ± standard deviation (SD) of at least two or three independent experiments. Data were statistically significant when P<0.05.

## Acknowledgments

We thank Stephanie Deming, Research Medical Library, MD Anderson Cancer Center, for editing the manuscript. This research was supported in part by the John S. Dunn Foundation, the John S. Dunn, Sr. Distinguished Chair in Diagnostic Imaging, the University Cancer Foundation via the Institutional Research Grant program at the University of Texas MD Anderson Cancer Center, NIH/NCI grant R01CA258540 (C.L.); NIH/NIAID grant 1R21AI101932 (Q.M.), NIH/NCI grant 1P01CA148600 (Q.M. and J.J.M.).

## DECLARATIONS

### Authors’ contributions

D.L., S.X.S., Q.M., and C.L. conceived and designed the study. D.L. and S.X.S., and Q.Z.C. performed the experiments and provided data. A. K. S. and J.J.M., provided important intellectual content. D.L., S.X.S, and C.L drafted the manuscript. All the authors discussed the results and commented on the manuscript.

